# Characterization of brown-black pigment isolated from soil bacteria, *Beijerinckia fluminensis*

**DOI:** 10.1101/2021.07.21.453201

**Authors:** Mahesh H. Joshi, Ashwini A. Patil, Ravindra V. Adivarekar

## Abstract

Melanin is a ubiquitous pigment found in most organisms, it is a dark-brown or black pigment formed by the oxidation of phenolic compounds. They are negatively charged amorphous compounds having quinone groups. In this study; melanin-producing microorganism was isolated from soil obtained from iron ore mine. The soil was enriched in modified Ashby’s glucose broth for 15 days at 30^°^C further to which it was isolated on modified Ashby’s agar at 30^°^C for seven days; the colonies showing pigmentation were selected for further study. Conditions were optimized for maximal production of melanin pigment. The effect of carbon, nitrogen, tyrosine, and metal salts on pigment production was studied. Alkaline conditions were used to extract the pigment from cells, further characterized by UV-VIS spectroscopy for λ-max. FTIR was done to identify the native functional groups, and XRD was performed to determine the melanin’s structure. TGA analysis was done to check its thermal stability. SEM was carried out to check the size and shape of the melanin pigment. The melanin pigment was also analyzed for UV protectant property which was studied by exposure of both melanized and non-melanized cells to UV light at 254nm.

## Introduction

Melanins are negatively charged, hydrophobic macromolecules of high molecular weight formed by the oxidative polymerization of phenolic or indolic compounds. The resulting pigments are usually brown or black, but other colors have also been observed, depending on the substrate used to induce melanization. Melanin polymers are remarkable in that they have a stable population of organic free radicals. Melanin is widely distributed in living organisms such as bacteria, fungi, plants, animals, and human beings: eumelanins, pheomelanins, allomelanins, and pyomelanins. Eumelanins are formed from quinones and free radicals. Pheomelanins are derived from tyrosine and cysteine. Allomelanins are synthesized from nitrogen-free precursors, and pyomelanins are derived from tyrosine’s catabolism via p-hydroxyphenylpyruvate homogentisic acid (HGA) [1].

Melanin is an enigmatic high-molecular-weight pigment that has been extensively studied and characterized as a negatively charged amorphous compound with quinone groups. Structurally, Melanins are polyanionic molecules that contain a stable population of organic free radicals. Melanin is a dense, rigid, and amorphous complex absorbing material with significant strength. Melanin absorbs electromagnetic waves (or light) throughout the entire ultraviolet (UV) and visible spectrum (VIS) and is therefore photo-protectant and photoconductive [2]. Melanin is an important virulence factor in several human pathogenic fungi due to its anti-oxidative, anti-phagocytic, thermostable, anti-radioactive, paramagnetic, and metal-binding properties. The pigment also offers protection as a cellular scavenger against free radicals, ROS (reactive oxygen species), drugs, oxidants, and xenobiotics [3]. It has been reported that some microorganisms synthesize melanin in response to stress conditions such as high temperature, starvation, or hyperosmotic media. Melanized cells are more resistant to the effects of certain antifungals. Melanin pigment offers resistance to extreme heat and cold to the organism producing it. Various functional groups present on the surface of melanin structure offer heavy metal binding bioabsorption property and can be used for bioremediation [4]. Melanin can store and release endogenous and exogenous compounds, such as zinc, calcium, iron, and protein. Some drugs and other organic and inorganic substances, such as ampicillin, have also shown this pigment’s affinity. Characteristically, Melanins are insoluble in water or common organic solvents, resistant to concentrated acids, and susceptible to bleaching by oxidizing agents [5]. This study focuses on the isolation of microorganisms from soil contaminated with metals. The aim is to isolate and characterize microorganisms producing melanin and test the UV protectant ability of melanin.

## Materials and Methods

### Sample collection, enrichment, isolation, and identification

The soil samples were collected in a sterile container from the Tumsar region in the Bhandara district of Maharashtra, India. For enrichment; 1gm of soil sample was inoculated in the 100ml of Ashby’s broth medium and incubated at 30^°^C for 15 days. After enrichment, the isolation was carried out on Ashby’s glucose agar and incubated at 30^°^C for 7 days. The culture showing pigmentation was selected and was reisolated on Ashby’s glucose agar and preserved for further optimization studies. The culture’s identification was based on cultural, morphological, and biochemical characteristics as per Bergey’s manual of systematic bacteriology. Nomenclature & Phylogenetic analysis of bacterial culture has been done using bacterial 16S rRNA gene sequencing (∼1300 bases).

### Optimization of growth parameters for maximum pigment production

#### Growth medium

The growth medium was modified with different concentrations of carbon and nitrogen sources, L-tyrosine, and co-factors like Ammonium molybdate tetrahydrate [(NH_4_)_6_Mo_7_O_24_.4H_2_O], Ferrous sulphate [FeSO_4_.7H_2_O], and Magnesium sulphate [MgSO_4_.7H_2_O] to check the increase in melanin pigment production. To evaluate the optimum carbon source, different concentrations (0.1, 0.2, 0.4, 0.6, 0.8, and 1.0 %) of glucose, fructose, and sucrose were taken. Different concentrations of Nitrogen (ammonium sulphate) and L-tyrosine were prepared (0.01, 0.02, 0.04, 0.06, 0.08, and 0.10 %). Similarly, different concentrations (0.01, 0.02, 0.04, 0.06, 0.08, and 0.10 %) of (NH_4_)_6_Mo_7_O_24_.4H_2_O and FeSO_4_.7H_2_O were also prepared.

### Effect of aeration, incubation time, pH, temperature, and sunlight

The isolated culture was challenged with different physical parameters, like aeration, where one set of cultures was kept in an orbital shaker at 120rpm. The other in static condition both were incubated for 30 days at RT. pH selected were 3, 4, 6, 7, 8, 10, 12, 13. The effect of different temperatures (20, 24, 28, 37, 40^°^C) was also studied on the culture for maximum growth and pigment production. The culture inoculum was adjusted to 0.1O.D, and the media flasks (100ml) were kept for a week, and the interpretation was made by measuring absorbance at 400 nm using a UV-Vis spectrophotometer. [6].

### Growth characteristics of Beijerinckia fluminensis

The organism’s growth curve and maximum pigment production were studied using Ashby’s glucose and modified Ashby’s glucose medium. Estimation was done after the third day of incubation for 30 days. The interpretation was made by measuring the cell density (OD) and pigment absorbance at a fixed wavelength of λ-max of 600 nm and 400 nm [6].

### Extraction of melanin pigment

The extraction of the pigment was carried out using standard alkali solutions like NaOH, KOH, NH_4_OH, Ca(OH)_2_, LiOH at varying concentrations of 0.1, 0.5, 1, 1.5, 2 N. The extraction was carried out from a thirty-day old culture plate grown on MAG agar. After extraction. The pigment was precipitated using double strength acid like HCl (2N) under slow magnetic stirring conditions. Further, the residue was transferred to a dialysis tube and kept in distilled water for 6 h with two distilled water changes. After dialysis, melanin was re-dissolved in alkali and precipitated; this was done to altogether remove the unwanted impurities like proteins and other fatty acids present.

### Characterization of melanin extracted from *Beijerinckia fluminensis*

Characterization of melanin is a difficult task due to its insoluble nature. Scanned literature has shown significant characterizations like UV-Spectroscopy, FTIR, XRD, TGA, and SEM. UV-visible spectroscopy of bacterial melanin was carried out by dissolving the dry pigment in 5ml of 0.1 N NaOH, and absorbance was recorded using UV-1800 Shimadzu spectrophotometer against NaOH as blank. The spectra were compared with the standard melanins (isolated from bacteria and cuttlefish, *Sepia Officinalis*). FTIR (FTIR-8400S Shimadzu) was performed to confirm native groups of melanin compared with the other natural melanin present in higher organisms. XRD analysis was carried using Lab X, XRD-6100 of Shimadzu, after drying the melanin at 200^°^C and converting it into a fine powder. TGA data were recorded in an aluminum pan using 10mg melanin samples using a TGA Shimadzu (DTG-60H, Japan) thermal analyzer under an atmosphere of nitrogen from 30^°^C to 500^°^C at a heating rate of 10^°^C/min. For SEM analysis, melanin was dried, and finely powdered melanin was mounted on an SEM stub (diameter, 12 mm×12mm) using carbon tabs. It was coated with Au/Pd in a sputter coater, and the specimen was then scanned in FEG-Philips XL 30 SEM.

### UV resistance property of bacterial melanin

Photo-protectant property of melanin was checked by initially growing the cells in two flasks a) Modified Ashby’s glucose broth (MAG) and b) MAG broth with L-ascorbic acid, which prevents the formation of melanin. These were incubated for 5 days on shaker condition at 25^°^C. On the 7^th^ day, 10ml of culture aliquots, each from both the sets (a and b), were taken in ten sterile empty Petri dishes under aseptic condition and exposed to UV radiation (254 nm) for different time intervals (0, 2, 4, 6, 8, 10, 15, 20, 25, 30 min) at a distance of 30cm from base to UV lamp.

Further, from each plate, 0.1ml of aliquot is added to 10ml MAG broth to check cell death at a particular time of UV exposure. At the same time, loopful was streaked on Modified Ashby’s glucose agar plates. Tubes and plates were incubated for 72h, at 25^°^C. The result’s interpretation was noted based on turbidity observed in the tubes and the recovery of growth on the line streaked on MAG agar plates.

## Results

### Enrichment, isolation, and characterization

#### Isolation and characterization

Isolation from the enriched broth on Ashby’s glucose agar showed large slimy colonies with pigmentation. These colonies were selected and reisolated on fresh agar plates. After growing extensively for 7 to 30 days, light brown to black pigmentation with slimy consistency was seen, which was maybe due to the production of polyhydroxbutarate (PHB) (Fig. 1). The growth biochemistry of *Beijerinckia* shows the utilization of glucose, fructose, and sucrose. The organism is catalase-positive with the presence of urease but absence of nitrate reduction capability. The optimal growth is seen between 20-30^°^C and between pH 3.0 and pH 9.0. When observed under the microscope, the cells appear straight or slightly curved rods with rounded ends. Cells occur singly or appear as dividing pairs. Intracellular granules of poly-b-hydroxybutyrate (PHB) are formed, generally one at each pole. Gram-negative, motile by peritrichous flagella or nonmotile [7,8].

**Fig. 1.**
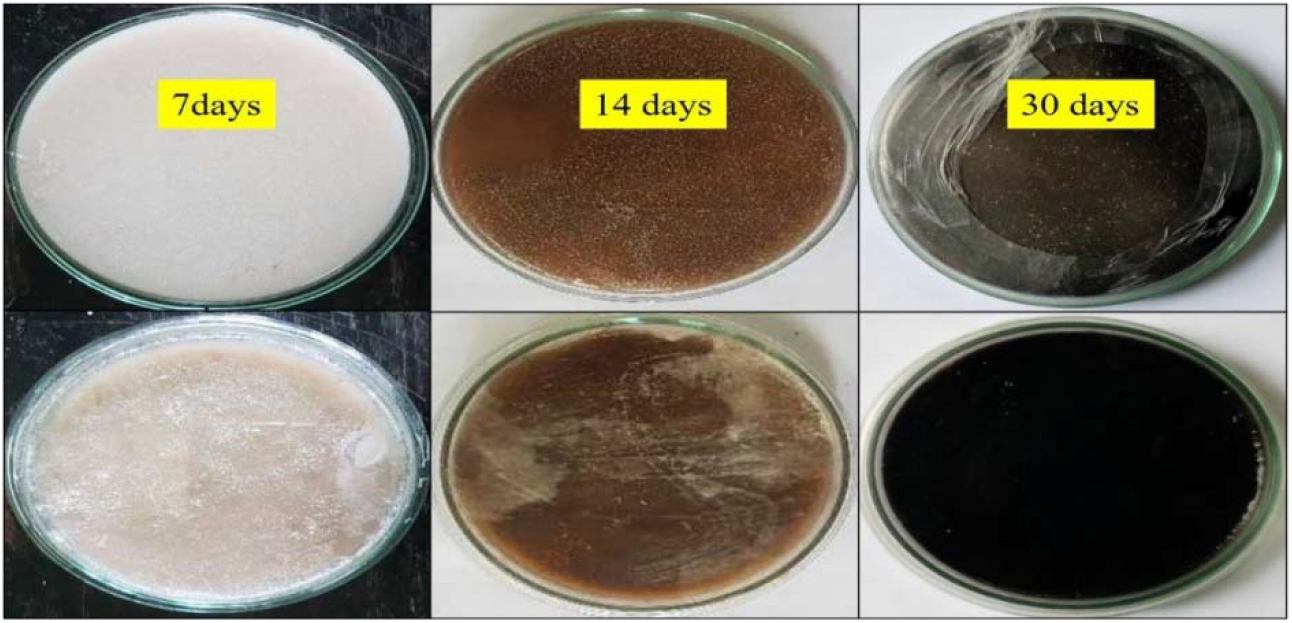
Ashby’s glucose agar plates showing melanin production and growth of isolated bacteria.

The culture’s physical and biochemical characteristics partially confirm the isolate to be from the *Azotobacter* spp. or *Beijerinckia* spp. (Table 1). Table 2 shows the 16S ribosomal RNA sequence, which confirms the *Beijerinckia* genus of the isolated strain, and the NCBI-BLAST analysis showed 100% homology with the *Beijerinckia fluminensis* strain UQM 1685^T^ [9].

**Table 1.** Physical and biochemical characteristics of a strain isolated from soil.

| Characteristics | Result |
| --- | --- |
| Colony | Irregular |
| Consistency | Slimy (due to presence of PHB) |
| Pigmentation | Light to dark brown |
| Motility | + |
| Gram nature | gram-negative |
| Nitrate to nitrite reduction | – |
| Urease | – |
| Glucose | + |
| Fructose | + |
| Sucrose | + |
| Mannitol | – |
| Maltose | – |
| Lactose | + |
| Glycerol | + |
| Sorbitol | + |
| Key: +, positive; –, negative |  |

**Table 2.** 16S ribosomal RNA gene sequence and NCBI-BLAST analysis with Phylogenetic tree of *B. fluminensis* showing homology with the species.

| Sr. No. | Strain | Aim | Sequence | NCBI BLAST | Remarks |
| --- | --- | --- | --- | --- | --- |
| 1. | Seq 34_AZM_<br>NC160318A | Identification<br>of 16S rRNA<br>gene | F27_1492R<br>C(1303bp) | NR_116306<br>*Beijerinckia fluminensis*<br>Identities:<br>1302/1303(99%)<br>Int. J. Syst. Evol.<br>Microbiol. 59 (PT 9),<br>2323-2328 (2009)<br>++<br>NR_115516<br>*Agrobacterium*<br>*tumefaciens*<br>Identities:<br>1296/1303(99%)<br>J. Gen. Appl. Microbiol.<br>49 (3), 155-179 (2003)<br>++<br>NR_116874<br>*Rhizobium pusense*<br>Identities:<br>1294/1304(99%)<br>Int. J. Syst. Evol.<br>Microbiol. 61 (PT 11),<br>2632-2639 (2011) | Strain shows<br>closer<br>homology with<br>*Beijerinckia* sp.<br>(Closer<br>to *fluminensis*) |

### Optimization studies

#### Optimization for maximum pigment production

Optimization of pigment production results showed the strain could utilize all three carbon sources and indicated that as the sugars’ concentration increases, the pigment production also increased exponentially. The culture showed a high melanin production rate while growing on glucose as the sole source of carbon (Fig. 2). The effect of L-tyrosine was significant on melanin production since it is one of the main precursors in the melanin pathway. The effect nitrogen source was not substantial, and the pigment production over the range was average (Figure 3). Figure 4 shows the effect of growth co-factors like ammonium molybdate tetrahydrate [(NH_4_)_6_Mo_7_O_24_.4H_2_O], ferrous sulphate [FeSO_4_.7H_2_O], and Magnesium sulphate [MgSO_4_.7H_2_O], which showed a positive growth response in maximizing pigment production from 0.04% to 0.06% concentration. *Beijerinckia* sp are aerobic and grow well in conditions having optimum oxygen levels. This study was carried out to analyze the effect of aerobic conditions on melanin production. Melanin production was significant in the culture flask kept in an orbital shaker, which showed melanin production in six days.

**Fig. 2.**
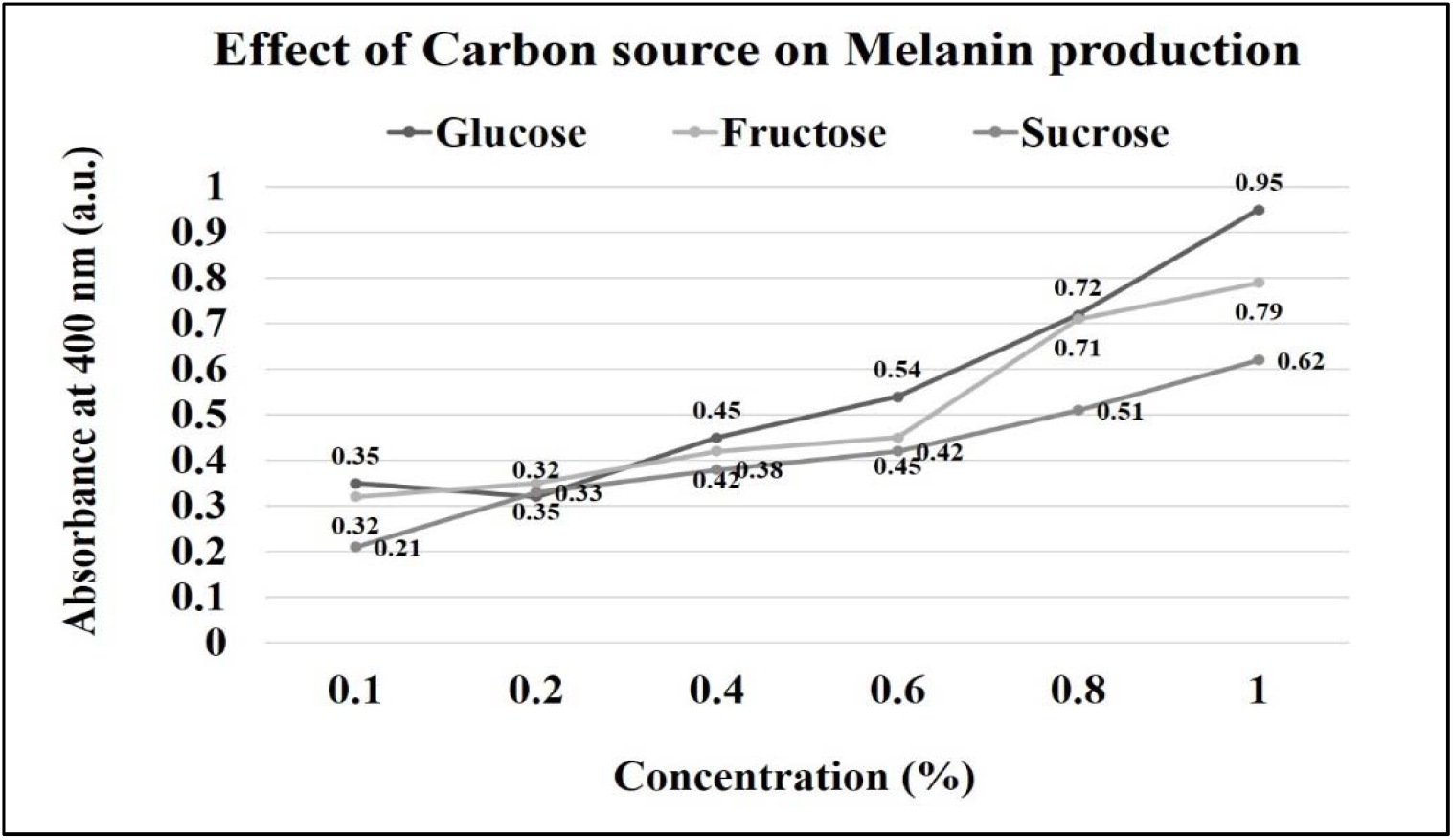
Effect of carbon source on melanin production

**Fig. 3.**
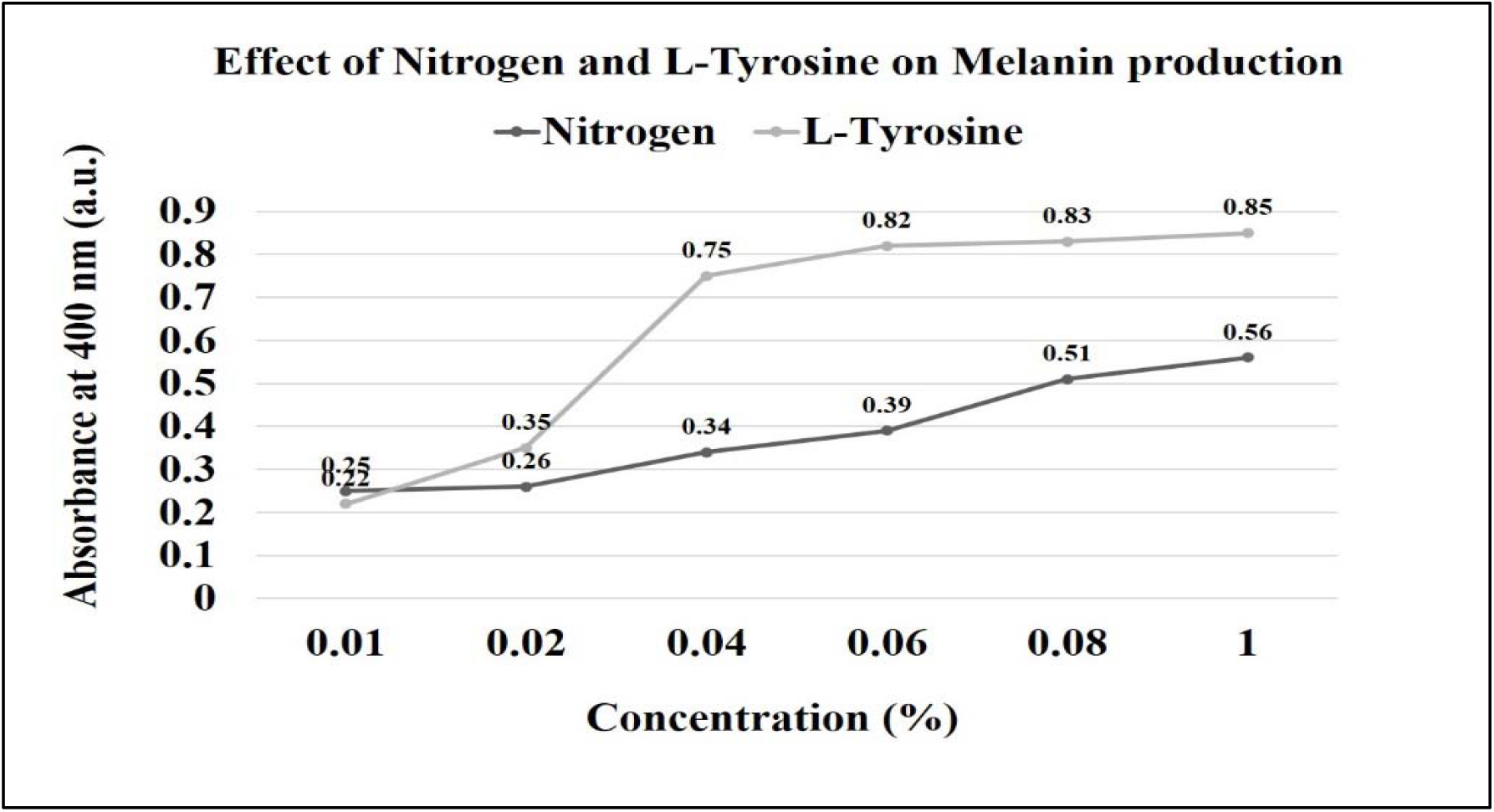
Effect of nitrogen and L-tyrosine on melanin production

**Fig. 4.**
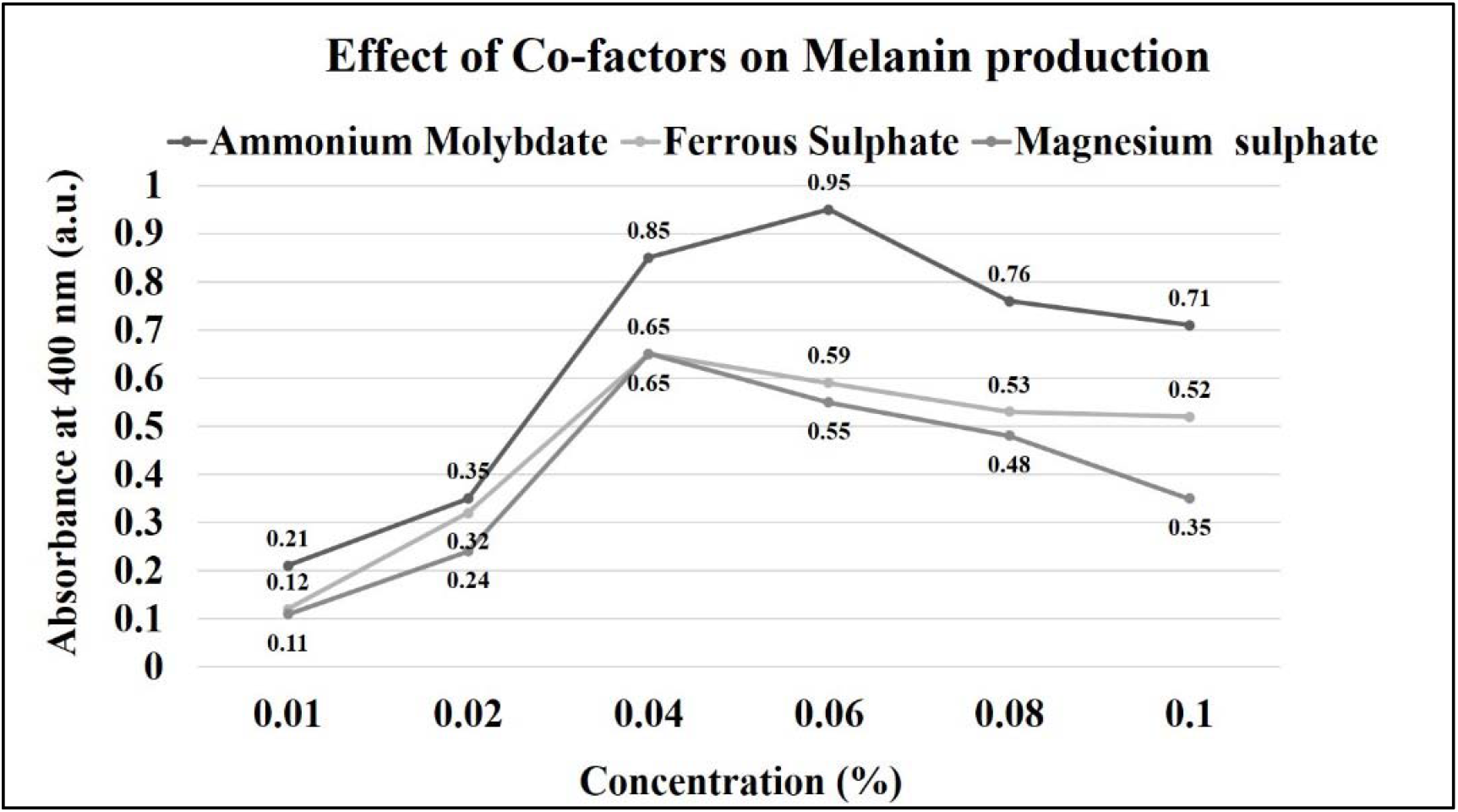
Effect of co-factors on melanin production

In contrast, the flask kept static showed melanin production after ten days. As the incubation time increased, the melanin production also increased, which was studied for thirty days. *Beijerinckia* cells grow slowly and require the right amount of oxygen for PHB production. In shaker conditions, the cells utilize growth nutrients faster and move early to the stationary phase, which results in the transition of older cells to stress conditions, leading to melanin production (Fig. 5) [10,11].

**Fig. 5.**
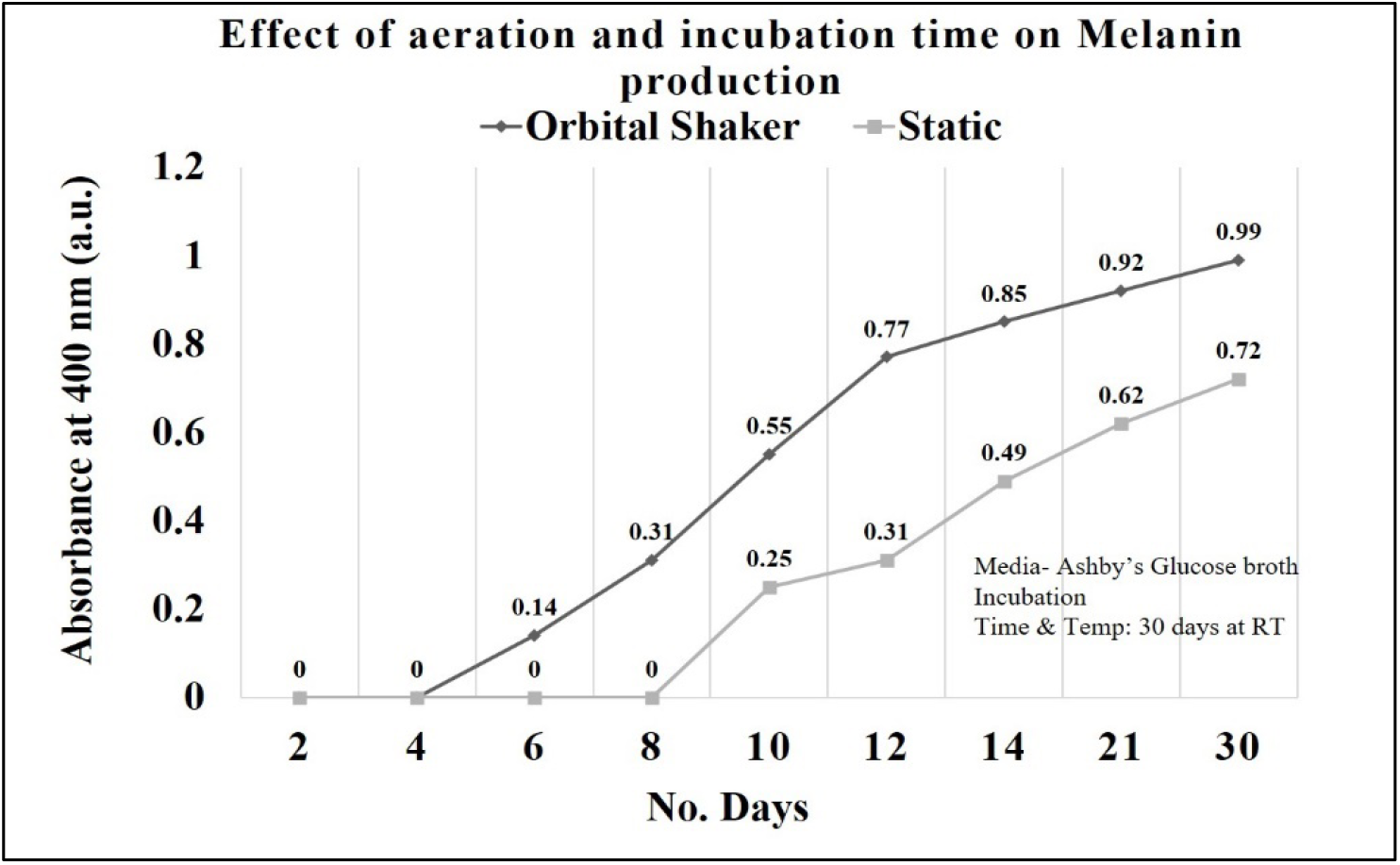
Effect of aeration and incubation time on melanin production in *B. fluminensis*

The pH of the medium and incubation temperature plays a vital role in the functioning of the cell’s metabolic activities. The earlier studies understand that the *Beijerinckia* sp. grows better in the pH range of 3.0 to 9.0 at an optimal temperature of 25^°^C to 30^°^C [13,14]. Keeping this information in mind, the effect of different pH and temperature was studied on melanin production, and it was found that the maximum output of melanin was seen from pH 4.0 to pH 6.0 and the optimal growth temperature for significant melanin production was 30^°^C (Fig. 6).

**Fig. 6.**
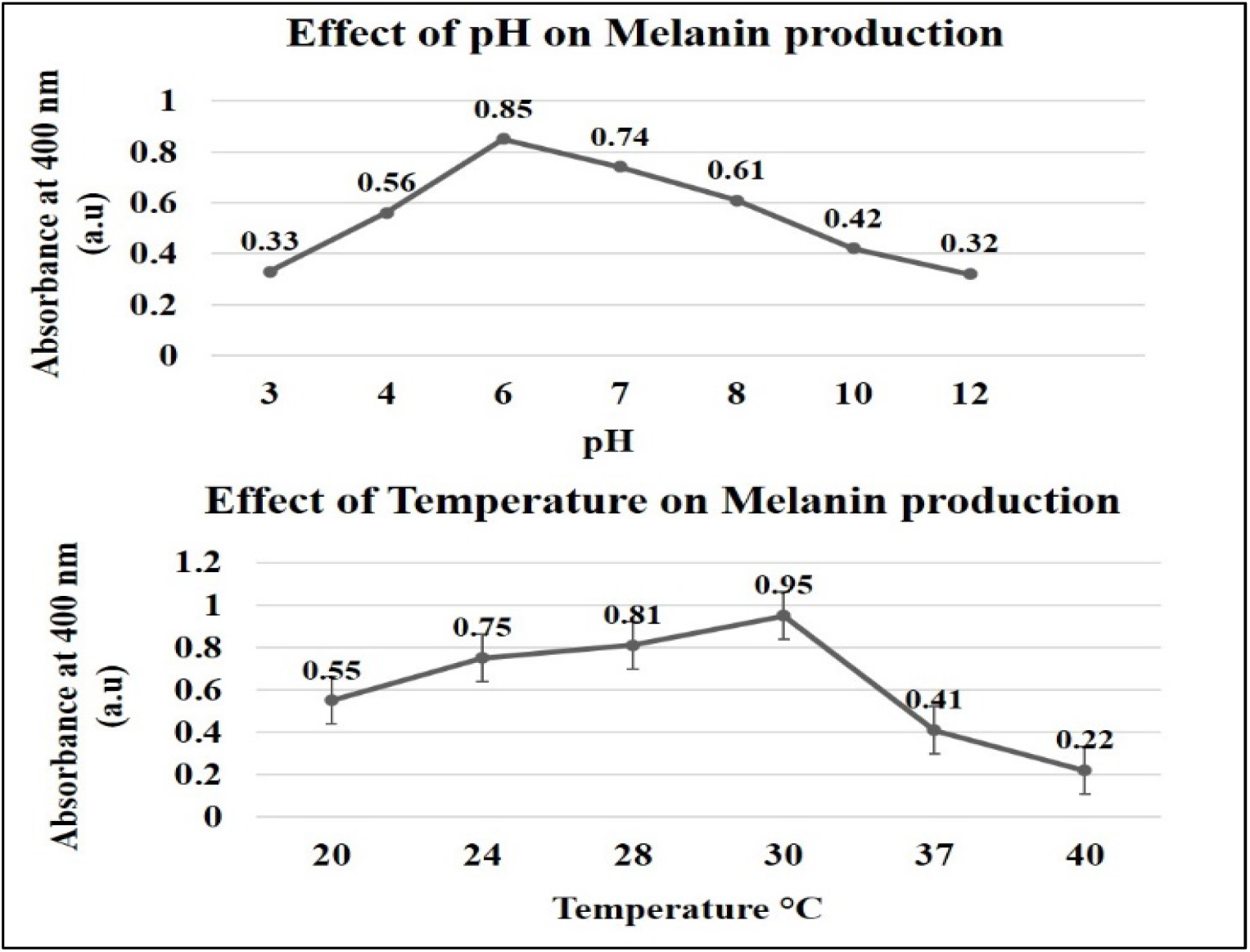
Effect of pH and temperature melanin production in *B. fluminensis*

#### Growth characteristics of Beijerinckia fluminensis

The growth characteristics and pigment production of *B. fluminensis* in Modified Ashby’s glucose medium were better than those in the non-modified medium. The presence of Nitrogen, L-tyrosine, and co-factors in the medium was found to enhance the growth and double the melanin production of the culture (Fig. 7).

**Fig. 7.**
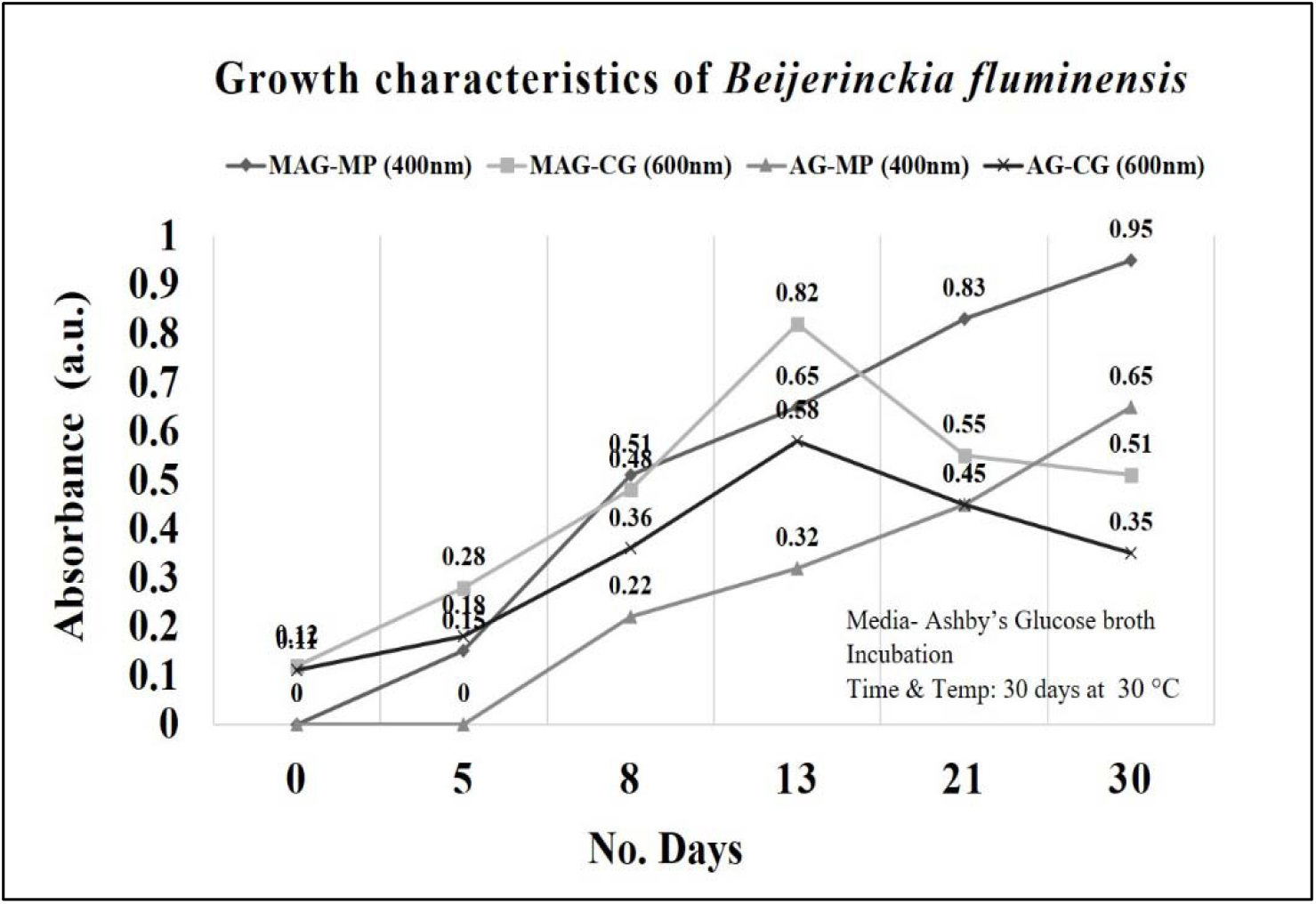
Growth characteristics of *B. fluminensis* in Ashby’s glucose (AG) and modified Ashby’s glucose (MAG)

### Extraction of melanin pigment from grown culture

From the literature, it is evident that the melanins are only soluble in alkaline conditions. The melanin pigment extraction was carried out from the isolate, grown on Ashby’s glucose agar for thirty days (Fig 8). 1gm of the solid media was transferred in tubes having 10 ml of different alkalis. The tubes were vortexed, and the solutions were kept in a sonication bath for 30 min. Based on the solution’s clarity, results were noted as either “soluble” or “insoluble.” The melanin showed better extractability in strong alkalis like sodium hydroxide and potassium hydroxide (Table 3).

**Table 3.**
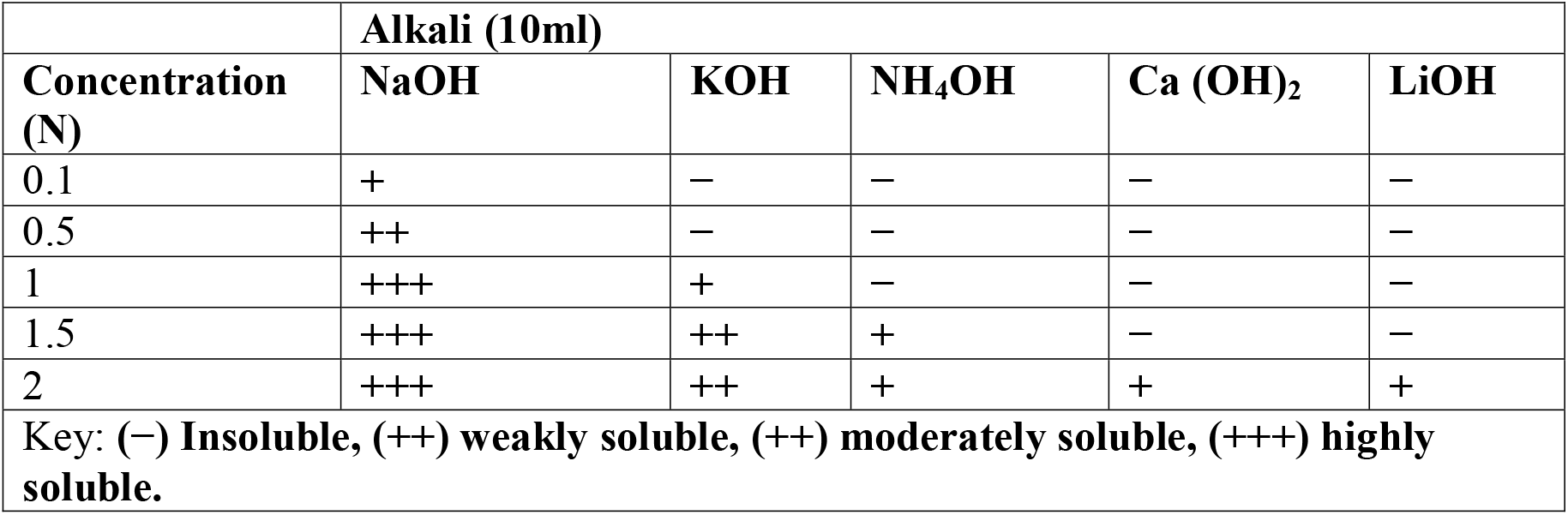
Extraction of melanin from *B. fluminensis* in different alkalis

**Fig. 8.**
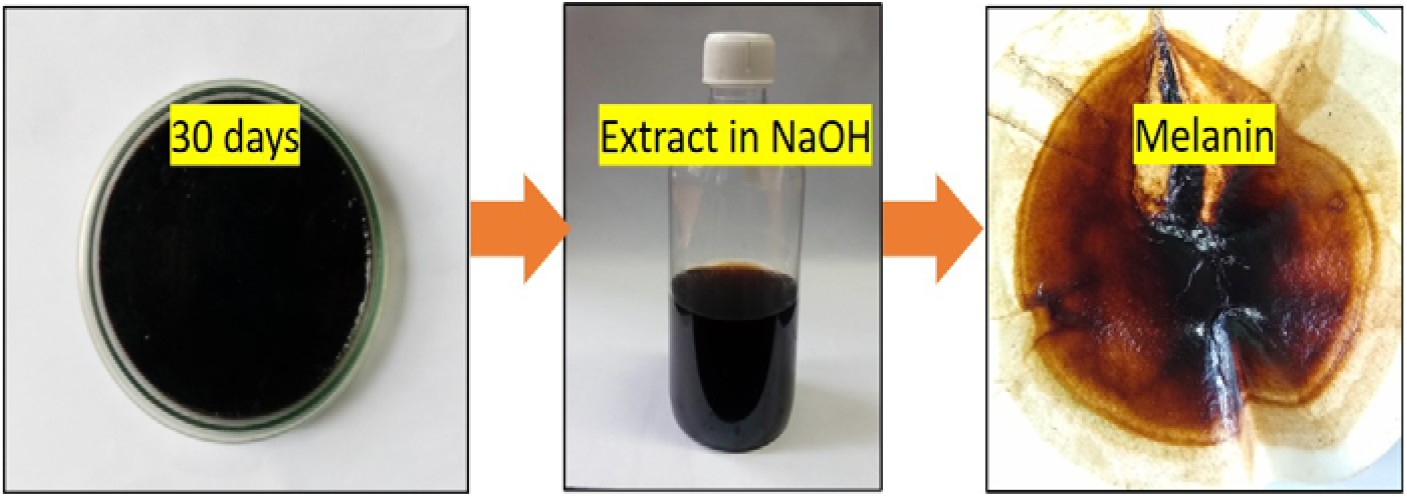
Extraction of Melanin from the thirty-day old grown culture of *B. fluminensis*

### Characterization of melanin from *Beijerinckia fluminensis*

The melanin pigment isolated from the bacteria was characterized by determining the λ-max using a 200-800 nm scanning range using a UV−Visible Spectrometer. The scan showed a peak in the UV region at 280 nm and was seen to decrease gradually. Mostly, the absorption was highest in the UV region at 200−300 nm (Fig. 9), which is known as the characteristic property of melanin [19,28]. The FTIR spectra (Fig.10) of melanin show a broad absorption peak at 3433 cm^-1^ and 3344 cm^-1^ which was due to the −OH and −NH2 groups’ stretching vibration. The peak observed at around 2857 cm^-1^ is assigned to C−H stretching vibration. The peak appearing in this range was typical of aromatic C−H groups from indole moieties in melanin pigment molecules. The absorption peak at 1676 cm^-1^ and 1618 cm^-1^ attributed to C−C or C−O stretching and that at 1460 cm^-1^ due to C−H bending mode were obtained from the spectra of other microbial melanin pigments [24,25].

**Fig. 9.**
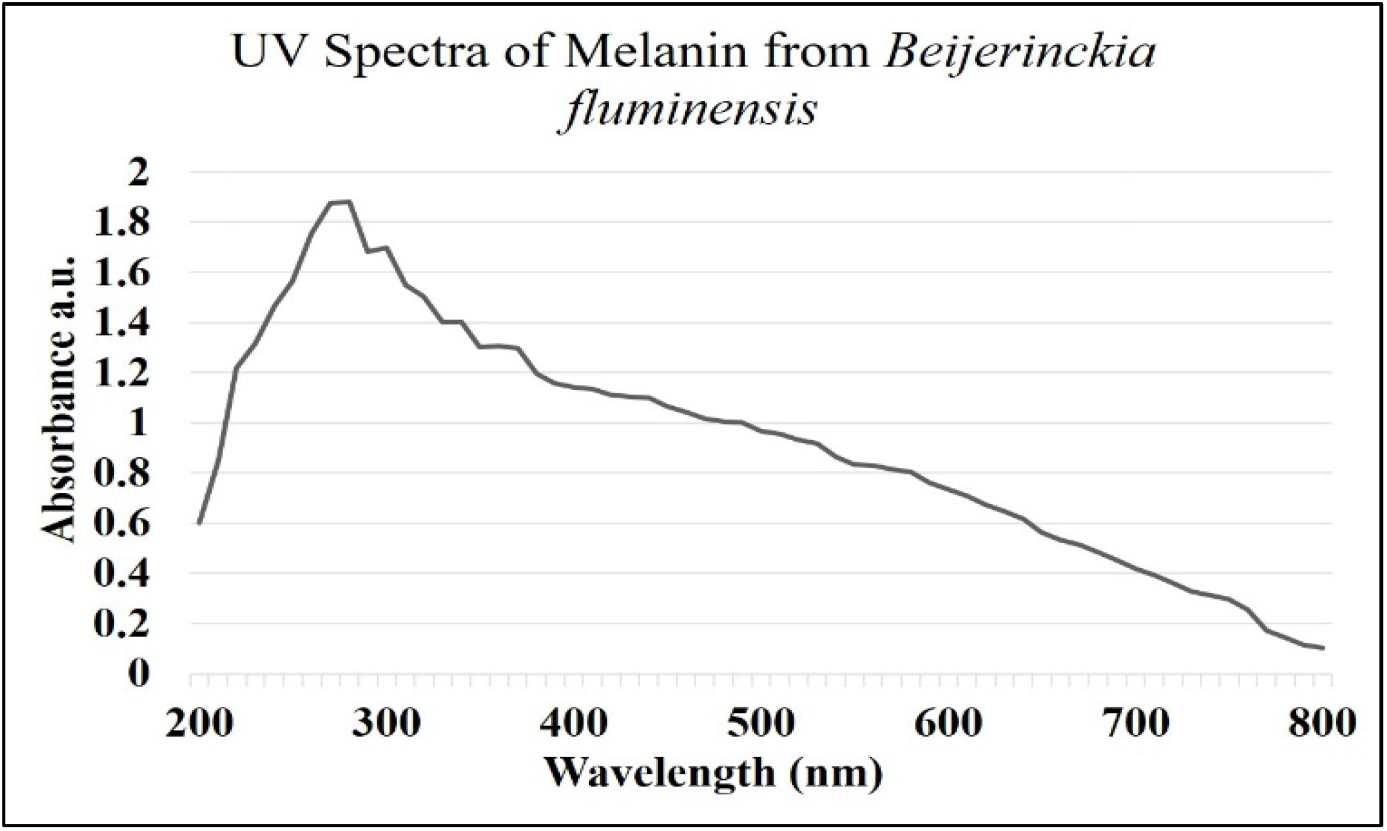
The UV-VIS Spectra of melanin from *B. fluminensis*

**Fig. 10.**
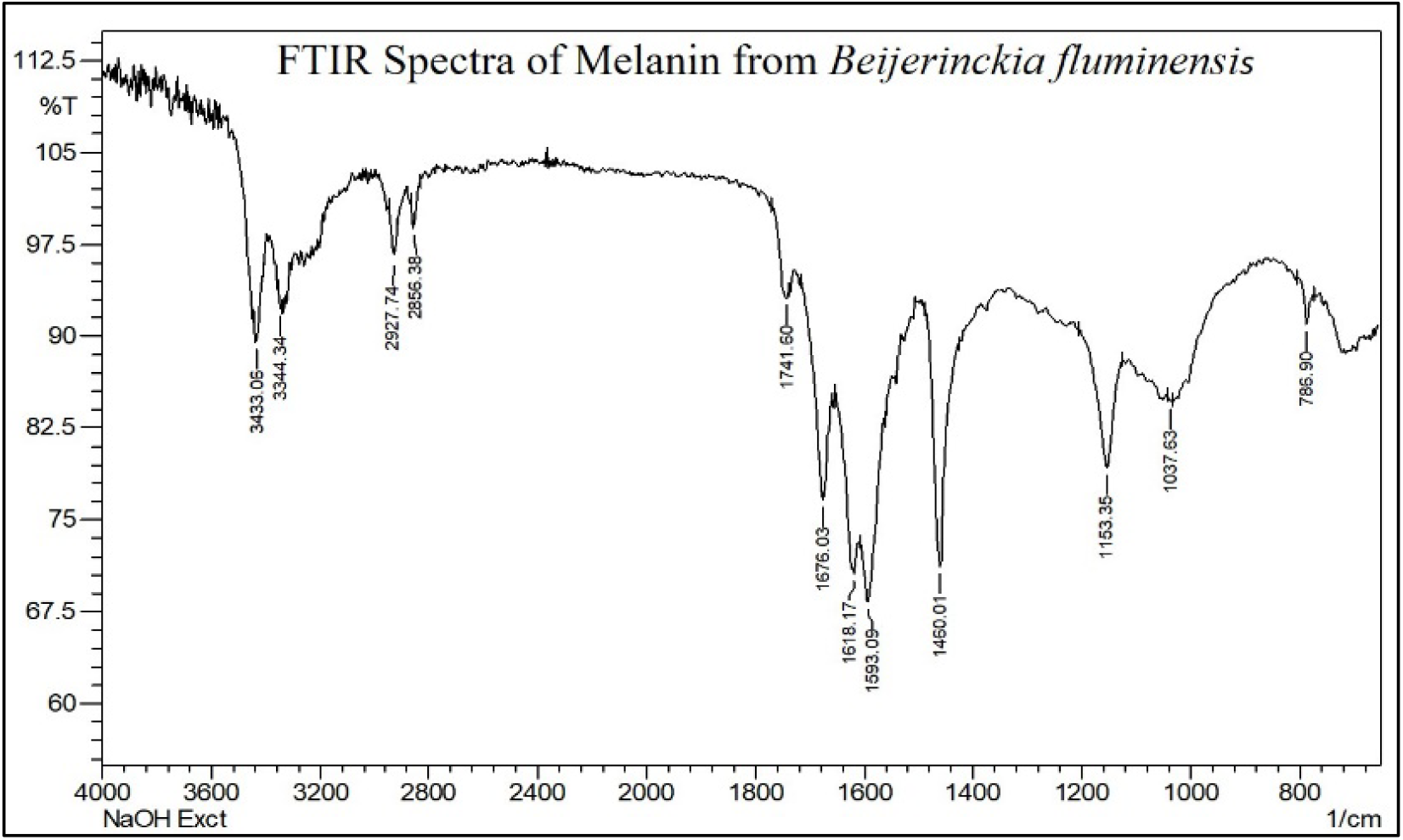
The FTIR spectra of melanin from *B. fluminensis*

The XRD spectrum (Fig. 11) of the dry melanin shows a broad diffraction peak (2θ=15-25^°^). Such broad peaks indicate non-Bragg features and present different orientations of the structural elements, which are random. Broad peaks are also characteristics of amorphous nature, which is seen in melanins. Melanin structures are uncertain [26,28] due to the amorphous, heterogeneous and insoluble nature of these pigments. TGA analysis of melanin show stability of the pigment at the higher temperatures of 500^°^C. Between 300^°^C and 400^°^C, the weight loss was 50%, a high percentage of which could be explained by considering melanin degradation because the purification processes removed all cellular molecules (Figure 12). A gradual weight loss was observed up to 500^°^C, indicating the complete combustion of melanin pigment. The SEM micrographs show irregular sizes of melanin particles (20µm to 2µm), which may be due to the melanin particles’ aggregation after drying (Fig 13). Melanins from different sources show different microstructures [29-32].

**Fig. 11.**
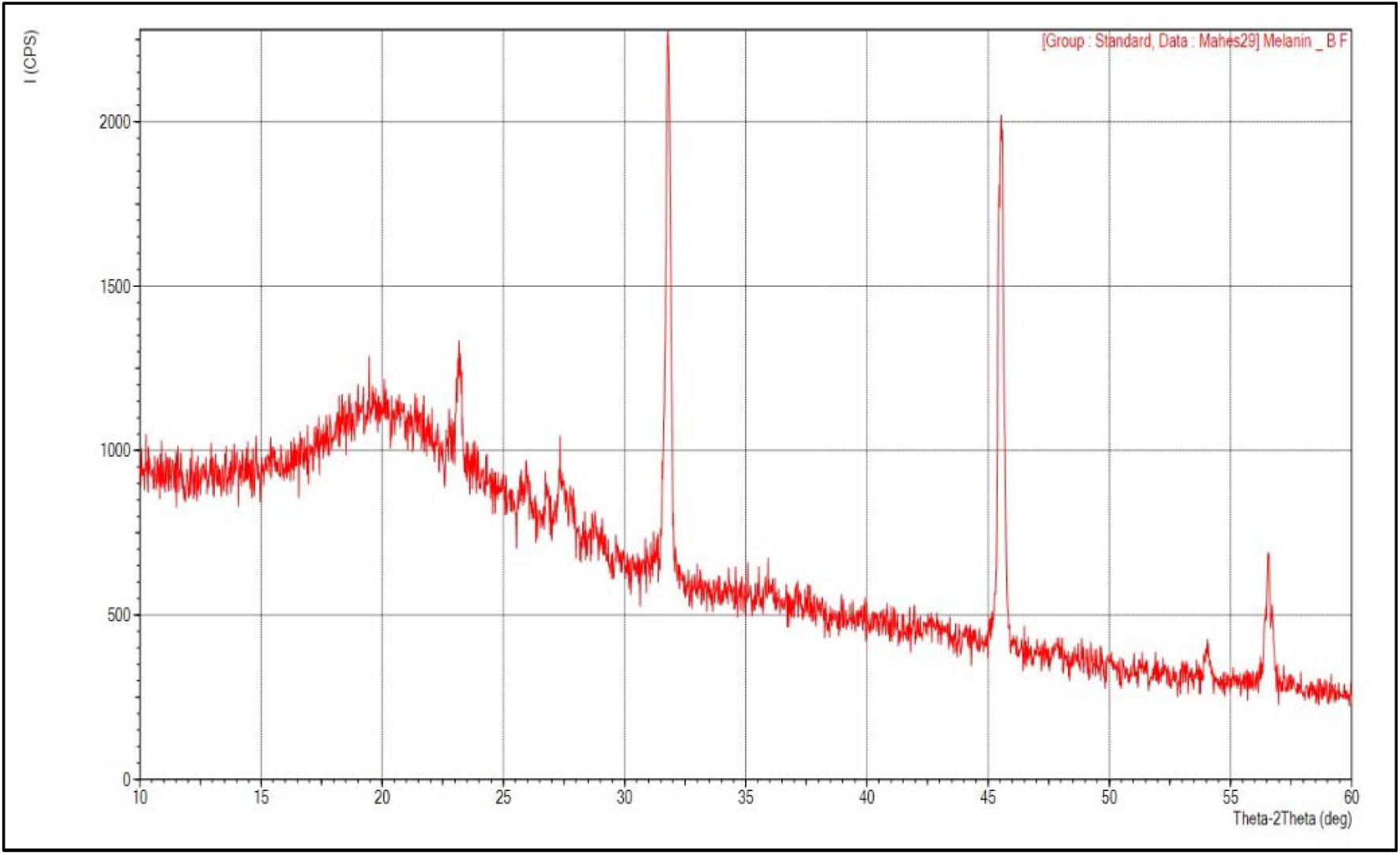
The XRD analysis of melanin from *B. fluminensis*

**Fig. 12.**
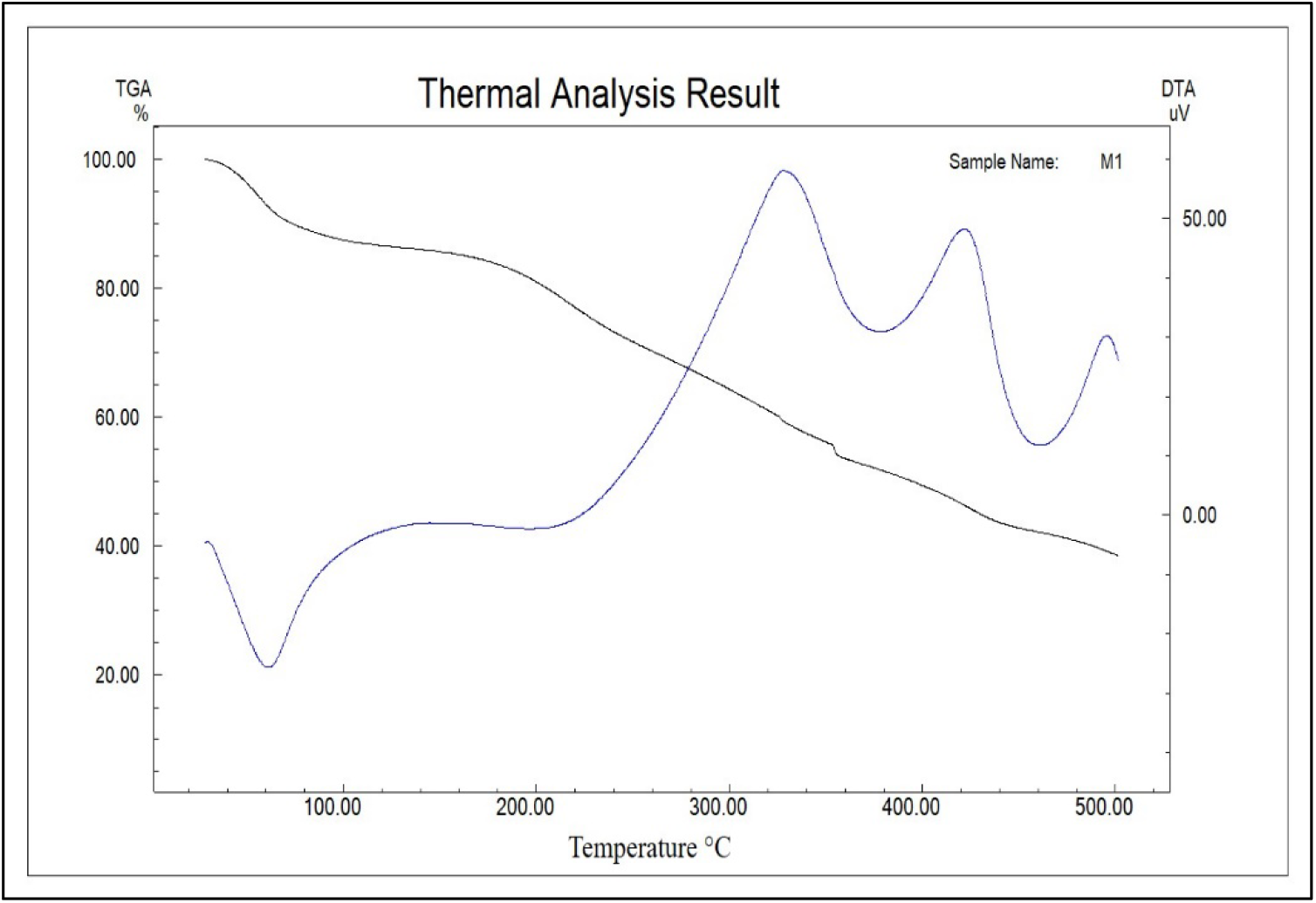
TGA thermogram of melanin from *B. fluminensis*

**Fig. 13.**
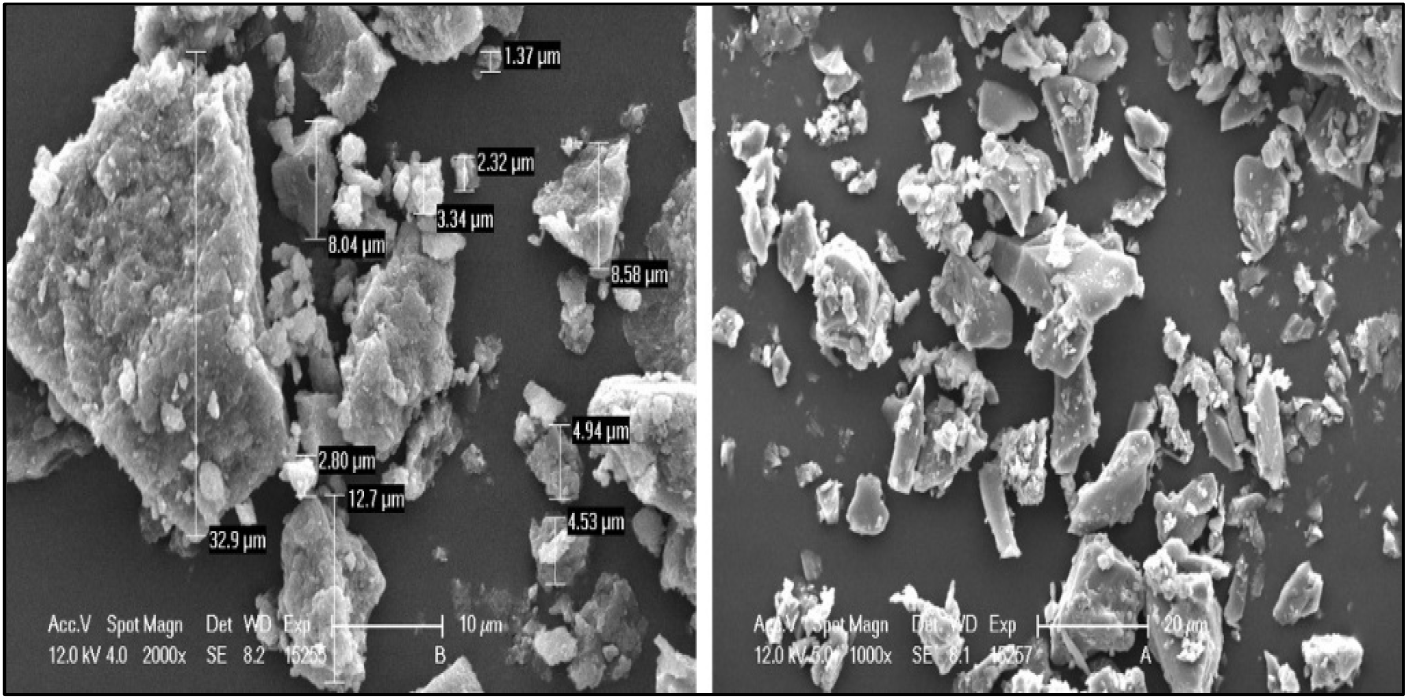
SEM analysis of melanin from *B. fluminensis* showing irregular sizes

**Fig. 14.**
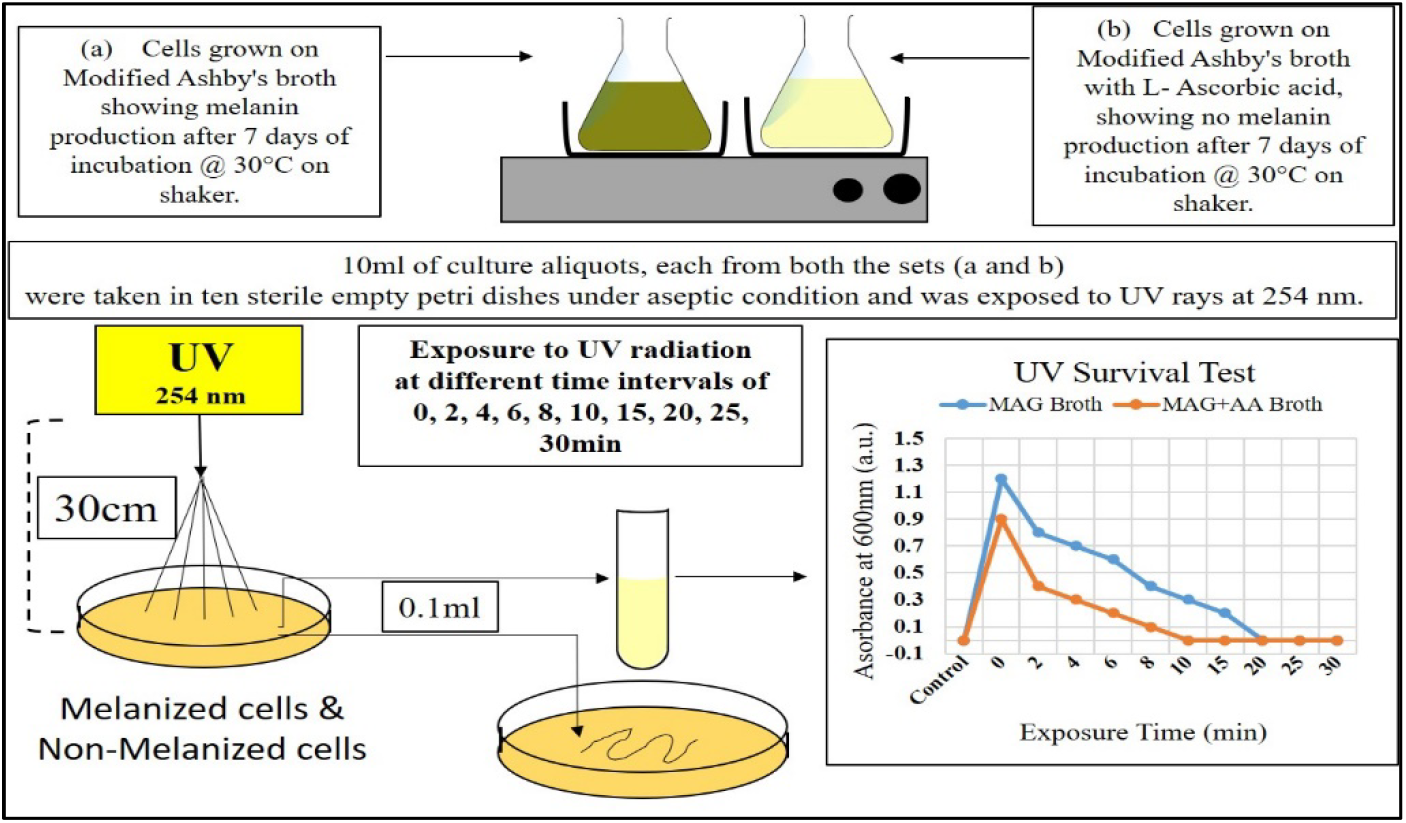
Schematic representation of UV the survival test of *B. fluminensis* cells exposed to UV radiation at 254nm.

### UV resistance property of melanin from *Beijerinckia fluminensis*

The UV-resistant property of bacterial melanin was tested using *Beijerinckia fluminensis* as the study model (Fig.14). The bacterial cell’s ability to produce melanin is suppressed using L-Ascorbic acid in modified Ashby’s broth. Both suppressed cells and normal cells were then exposed to UV radiation (254 nm) and showed variable results which are presented in table 4. The normal cells producing melanin showed no growth in 20 min of UV exposure, which was confirmed by the broth’s absence of growth and on plates incubated for more than seven days. The suppressed cells showed senescence in 8min of UV exposure time, where no growth was observed in further incubation.

**Table 4.** Evaluation of UV Protection property of melanin produced by *B. fluminensis*

| Exposure Time (min) | Modified Ashby's glucose broth |  | Modified Ashby's glucose broth & L-ascorbic acid |  |
| --- | --- | --- | --- | --- |
|  | Broth | Plate | Broth | Plate |
| Media Control | — | — | — | — |
| 0 | + | + | + | + |
| 2 | + | + | + | + |
| 4 | + | + | + | + |
| 6 | + | + | + | + |
| 8 | + | + | — | — |
| 10 | + | + | — | — |
| 15 | + | + | — | — |
| 20 | — | — | — | — |
| 25 | — | — | — | — |
| 30 | — | — | — | — |

Key: (+) Growth (—) No growth

## Discussion

The present study provides data for melanin pigment production from a non-pathogenic simple soil bacterium. Lasker et al. 2010, found *Beijerinckia* sp. in soil samples while studying rice plants’ rhizosphere [7]. It was observed that the optimum conditions like temperature and pH required for maximal pigment production from *B. fluminensis* were found significant. Oggerin et al. 2009, had studied the 16S rRNA gene sequence of *B. fluminensis* strains UQM 1685^T^ and CIP 106281^T^ and found that it was 90-91% similar to the other *Beijerinckia* species and subspecies [9]. Barbosa et al. [10] observed an increase in slime production and cell density in *B. derxii* over the time when glucose was the sole carbon source. Barbosa et al. 2002, have also shown that co-factors like metal salts and aerobic conditions are essential for the cell growth and specific nitrogenase activity of *B. derxii* strain ICB-10 [11]. Becking, observed that *Beijerinckia* species grow in the pH range of 3.0 to 9.0 at an optimal temperature of 25^°^C to 30^°^C [12,13,14]. They also require iron and molybdenum for optimal growth and N_2_ fixation; the molybdenum requirement was in the range 4.0−35.0 mg/l. The extraction of melanin from sources like hairs, cuttlefish, or microorganisms is extracted using either acid or bases. Generally, it is seen that melanins have good solubility in mild to strong bases like sodium and potassium hydroxide [15,29]. Bacterial melanin extraction is generally challenging due to cell membrane resistance; hence pretreatment with acids like HCL is necessary [15]. Extraction of fungal melanin from *Boltus griseus* was carried out using 1M NaOH with the treatment of 6M HCL [16,17]. Similar extractions were observed for microorganisms like *C. neoformans* and *C. sphaerospermum* [18].

Melanin has unique UV absorption spectra; first, sharp peaks are seen between 200−300nm, and the further the curve moves downwards monotonically. Aghajanyan et al. 2011, have seen a similar UV absorption trend in bacterial melanin [19]. These absorption characteristics were consistent with the UV-visible absorption spectra of the melanins from sources like fungi, bacteria, and melanin from sepia officinalis [20,21,22]. The FTIR spectra of melanins show bands that appear in the aromatic region having −C–H groups, carbonyl −C=O, a carboxylic acid group −COOH and −NH groups which are the main functional groups seen in melanins. Selvakumar et al. 2008, Araujo et al. 2012, and Joshi et al. 2019, had observed similar FTIR absorption peaks while studying melanins from a *Pleurotus cystidiosus*, marine sponges, and human hairs. The XRD spectra show a broad peak which is characteristic of melanin showing amorphous nature due to the non-Bragg features [26,28]. Thermogravimetric analysis of bacterial melanin showed higher temperature stability and is similar to that of a previously reported melanin [27,28]. Bacterial melanin isolated from *Streptomyces glaucenscens* shows porous and irregular structure [29,30]. Melanin from *Proteus mirabilis* is rounded aggregates of spherical bodies [31]. Similarly, *Pseudomonas sp*. and *Escherichia coli* produced melanin is found to be amorphous deposits with no differentiable structures and small granules [32].

L-ascorbic acid has a reducing effect on melanins’ precursors, e.g., it reduces the o-dopaquinone back to dopa, thus avoiding dopachrome and melanin formation [33,34]. Melanin-producing cells have a higher survival rate compared to non-melanized cells against UV radiation. The primary effect of UV rays is that it attacks the cell’s DNA as it contains high-energy photons. It leads to the formation of various mutations in (i.e., pyrimidine dimers, cross-linking of DNA and protein) the cell and undergoes senescence in the early growth phase. Other than the effect on DNA, the UV rays also have a role in producing reactive oxygen-derived free radicals, which directly impact the cell’s metabolic function. Thus, melanin plays a vital role in absorbing UV radiation and possibly prevents cellular damage [35,36].

## Conclusion

The present study provides data for melanin pigment production from a non-pathogenic soil bacterium. The biochemical and genetic analysis of the isolate confirmed the genus and species of the bacterium to *Beijerinckia fluminensis*. Optimum conditions required for maximal pigment production from *Beijerinckia fluminensis* was found significant. The physical parameters and shake flask data can be useful to carry out scale up for bulk production. Photo-protectant property of the melanin pigment were studied and thus confirms that melanin protects *Beijerinckia fluminensis* from harmful environmental conditions.

